# Chemogenetic manipulation of the ventral pallidum-nucleus accumbens shell pathway modulates sucrose consumption independent of motivation in female rats

**DOI:** 10.1101/2024.11.05.622115

**Authors:** Markie Peroutka, Ignacio Rivero Covelo

## Abstract

Here we investigated the role of the ventral pallidum (VP) to nucleus accumbens shell (AcbSh) pathway in mediating sucrose consumption and motivation for food in female rats. Using chemogenetic techniques, excitatory and inhibitory designer receptors were expressed in VP neurons projecting to the AcbSh. Behavioral assays assessed both lever pressing under a progressive ratio schedule and free sucrose consumption following activation or inhibition of this pathway. Results showed that inhibition of VP-AcbSh projections increased sucrose intake, while excitation reduced it, without affecting motivation to work for food. These findings indicate a specific influence of the VP-AcbSh pathway on sucrose consumption independent of food-seeking motivation. The study highlights the importance of differentiating VP efferents in understanding distinct roles in feeding behaviors and suggests that the VP-AcbSh pathway modulates consumption based on food type and potentially other variables.

## Introduction

The neural control of food intake and energy balance involves interactions between homeostatic and non-homeostatic systems. Traditionally, homeostatic regulation was attributed to hypothalamic and brainstem circuits responding to metabolic signals (Williams & Elmquist, 2012).

Critically, ventral striatopallidal structures, including the shell of the nucleus accumbens (AcbSh) and ventral pallidum (VP), exert a major influence on ingestive behavior by acting on some of these structures, mainly the lateral hypothalamus (LH). Inhibition of AcbSh neurons through GABA agonists or glutamate antagonists elicits intense feeding responses and activates LH neurons, as evidenced by increased Fos expression (Stratford & Kelley, 1999). The AcbSh projects to both the LH and VP, with unilateral lesions of either structure attenuating AcbSh-induced feeding (Stratford & Wirtshafter, 2012). The LH also modulates AcbSh activity directly through neurotransmitters like orexin and MCH, and indirectly via subcortical relay regions such as the VP (Castro et al., 2015; Urstadt et al., 2015). Relatedly, blockage of GABAA receptors in the VP elicits food intake in satiated rats (Stratford and Kelley, 1999), and this feeding presents a clear fat preference (Covelo et al., 2014).

Recent studies have suggested a role of sex in the mediation of sucrose consumption, as optogenetic stimulation of AcbSh projections to the VP decreases sucrose intake and alters hedonic value in female rats (Chometton et al., 2020). Parallelly, increases in sucrose intake have been reported in male, but not female rats, because of chemogenetic activation of GABAergic projection neurons in the VP (Scott et al., 2024).

Both the AcbSh and the VP regulate food intake. Notably, the relationship between the VP and AcbSh is that of a loop, and the role that projections between the two play in feeding remains understudied. The directionality of the circuit is relevant as projections from the AcbSh to the VP have different effects from projections from the VP to the AcbSh (Smedley et al., 2020). Additionally, as mentioned above, sex differences have been reported when modulating the projections of the VP (Scott et al., 2024). Here we aim to study the role that chemogenetic activation or inhibition of projections from the VP to the AcbSh have on the motivation to work for sucrose and on the consumption of sucrose in female rats.

## Materials and methods

### Subjects

250 to 300 g (time of arrival), female Sprague-Dawley rats (Envigo, Indianapolis, IN, USA) were used for these experiments. All rats were pair-housed in temperature and humidity-controlled rooms with a 12:12 light/dark cycle. In their home cages, rat pairs had access to chewing bones and a PVC pipe hut. After arrival at the facility, rats were allowed to acclimate to the colony room for at least 1 week before starting behavioral testing; during this time, the rats were handled once a day by researchers. Rats were also handled regularly for the duration of the behavioral experiments. All rats had ad libitum access to food and water for the duration of the experiments (except as noted below). Behavioral testing took place during the light cycle between 10:00 a.m. and 5:00 p.m. The experimental procedures were approved by the Institutional Animal Care and Use Committee at the University of Wisconsin-Parkside and were in accordance with the guidelines on animal care and use of the National Institutes of Health.

### Surgeries

Surgeries were performed using standard, aseptic, flat-skull stereotaxic techniques under isoflurane anesthesia (5% induction, 2% maintenance) delivered by a precision vaporizer. Once a stable plane of anesthesia was achieved, a sterile eye ointment was applied to both eyes (to prevent corneal desiccation), the analgesic administered, the scalp prepped for an incision (hair trimming, alcohol and iodine scrub), the incision exposed the skull and burr holes were performed above the target structures for the injection of adeno-associated viruses (AAV).

An AAV DIO construct containing an inverted form of either Gi (AAV5 AAV-hSyn-DIO-hM4D(Gi)-mCherry, Addgene Watertown, MA, USA) or Gq (AAV5 AAV-hSyn-DIO-hM3D(Gq)-mCherry, Addgene Watertown, MA, USA) DREADD was injected into the VP (from bregma: anterior posterior: -0.2 mm; medial lateral: ±1.8 mm; dorsal ventral: -8.7 mm). A retrograde AAV-Cre viral vector (AAVrg pENN.AAV.hSyn.HI.eGFP-Cre.WPRE.SV40, Addgene Watertown, MA, USA) was injected into the AcbSh (from bregma: anterior posterior: 1.6 mm; medial lateral: ±0.8 mm; dorsal ventral: -8.1 mm). Injections were performed using a Harvard micropump, Hamilton micro syringes connected to fluid-filled flexible tubing, and Plastics One injectors for a final volume of 1ml at an injection rate of 300 nl per minute.

For pain management, meloxicam (2 mg/kg, subcutaneous) was administered during the surgery and 24 hours later. Triple antibiotic was applied around the incision after closure using wound clips. Clips were removed 7 to 10 days after the surgery. Rats were allowed to recover for two weeks before behavioral testing.

### CNO preparation

Clozapine-N-oxide (CNO) was obtained from the National Institute of Mental Health (NIMH) Drug Supply Program. CNO was administered intraperitoneally 20 minutes before behavioral testing at a dose of 3.0 mg/kg. CNO was freshly prepared daily by dissolving it in 100% dimethyl sulfoxide (DMSO) and then diluting it with sterile water to a final concentration of 6% DMSO. A 6% DMSO solution in sterile water was used as the vehicle (VEH) control.

### Progressive ratio operant task

Rats were trained in a progressive ratio (PR) operant task using identical standard twin lever operant chambers (Med-Associates, St. Albans, VT) housed within sound attenuating chambers. First, animals got two daily, 30-min magazine training sessions in the operant boxes during which reinforcers (45 mg Precision Dustless banana pellets, BioServe, Frenchtown, NJ) were presented at 1-min intervals, with a “click” generated at the same time as food delivery. Next, rats were shaped to lever press and then placed on a fixed ratio (FR) 1 reinforcement schedule for two days. Rats got one session of training on an FR2 schedule followed the next day by one on an FR4 schedule. Subjects were then switched to a PR6 schedule, which continued for the remainder of the experiment. Each day, rats were placed into operant chambers with the house light on and both levers extended; only one lever was associated with food reward, although presses on both levers were recorded. The first response on the correct lever was followed by food reward, paired with the operation of the clicker. The number of responses required to earn each subsequent food pellet was increased by six after each reinforcer, so that seven responses were required to earn the second pellet, 13 to earn the third, and so on. The time of each lever press was recorded.

Each session continued until a pause in responding of 3 min duration occurs, a cut-oM value which has been used in other studies (Reilly and Trifunovic, 1999, Covelo et al., 2012), or 60 min elapsed, at which time the house lights were turned oM, the levers retracted, and the rats removed from the chambers. The breaking point was calculated as the final ratio completed in the session. Animals ran for five days on the PR6 schedule prior to drug treatment. After that, and 20 minutes before behavioral testing, rats were injected with either the CNO (3.0 mg/kg) or vehicle. All rats were administered two injections of CNO on two different days and two injections of vehicle, also on two different days.

### Sucrose consumption test

Rats were placed in individual home cages with wired bottoms and given access to a 20% sucrose solution for 60 minutes. This procedure was repeated in two consecutive days to acclimate the rats to the sucrose solution and minimize neophobia. After these two days, rats were administered with either CNO (3.0 mg/kg) or vehicle 20 minutes before being placed in the individual home cages. The sucrose bottles were weighed before and after the experiment to measure consumption. As described before, all rats got two CNO and two VEH injections, each injection on a different day.

### Perfusion and tissue processing

After completing the behavioral experiments, rats were anesthetized with 5% isoflurane and transcardially perfused with 0.9% saline followed by 4% formaldehyde (pH= 7.4) for fixation. Brains were extracted, post-fixed in 4% formaldehyde for 24 hours at 4°C, and immersed in increasing concentrations of sucrose solutions every 24 h (10%, 20%, then 30% sucrose in 0.1M PBS, pH= 7.4) at 4°C over the course of 3 days. Brains were then encased in Tissue-Plus O.C.T. (Fisher HealthCare, Houston, TX), frozen using dry ice, and subsequently sectioned in the coronal plane (45 μm) using a cryostat.

### Immunohistochemistry

The accuracy of DREADD expression in the VP and AcbSh was assessed using immunohistochemistry aimed at visualizing mCherry protein in DREADD-expressing neurons using procedures described before (Campus et al., 2019). Free-floating coronal sections from the VP and AcbSh were first rinsed 3 times in 0.1M PBS (pH= 7.4).

Endogenous peroxidase activity was blocked by incubating sections in 1% H2O2 for 10 minutes, followed by 3 additional rinses. To prevent nonspecific binding of the secondary antibody, sections were incubated in 0.1M PBS containing 0.4% Triton X-100 (TX) and 2.5% Normal Donkey Serum (NDS) (Jackson ImmunoResearch Laboratories, Inc., West Grove, PA). Sections were then incubated overnight at room temperature in primary antibody (rabbit anti-mCherry, Abcam, Cambridge, UK, diluted 1: 30,000) in 0.1M PBS + 0.4% TX + 1% NDS. Then, sections were rinsed again before being incubated for 1 h in a biotinylated donkey anti-rabbit secondary antibody (Jackson Immunoresearch, West Grove, PA, diluted 1: 500) in 0.1M PBS + 0.4% TX + 1% NDS. Peroxidase staining was obtained by using a standard avidin-biotin procedure using the Vectastain Elite ABC Kit (Vector Laboratories, Inc., Burlingame, CA, diluted 1: 1000 for A and B). Chromogenic reaction occurred by incubating sections in a 0.1M PBS solution containing 0.02% 3,3’-diaminobenzidine tetrahydrochloride (DAB) and 0.012% H2O2. Sections were rinsed and stored at 4°C until mounted, air-dried, and cover-slipped using a toluene-based mounting medium (Permount, Thermo-Fisher Scientific, Waltham, MA). Bright-field images containing the VP or the AcbSh were captured using a Zeiss Axioscan light microscope and were analyzed by an experimenter blind to the experimental groups. The location of mCherry expression was confirmed using a rat brain atlas (Paxinos and Watson, 2013). The final number of rats per group were: Gi →11, Gq →10, and No DREADD → 12. A schematic representation of the approach and representative mCherry pictures can be found in figure 1.

**Figure 1.**
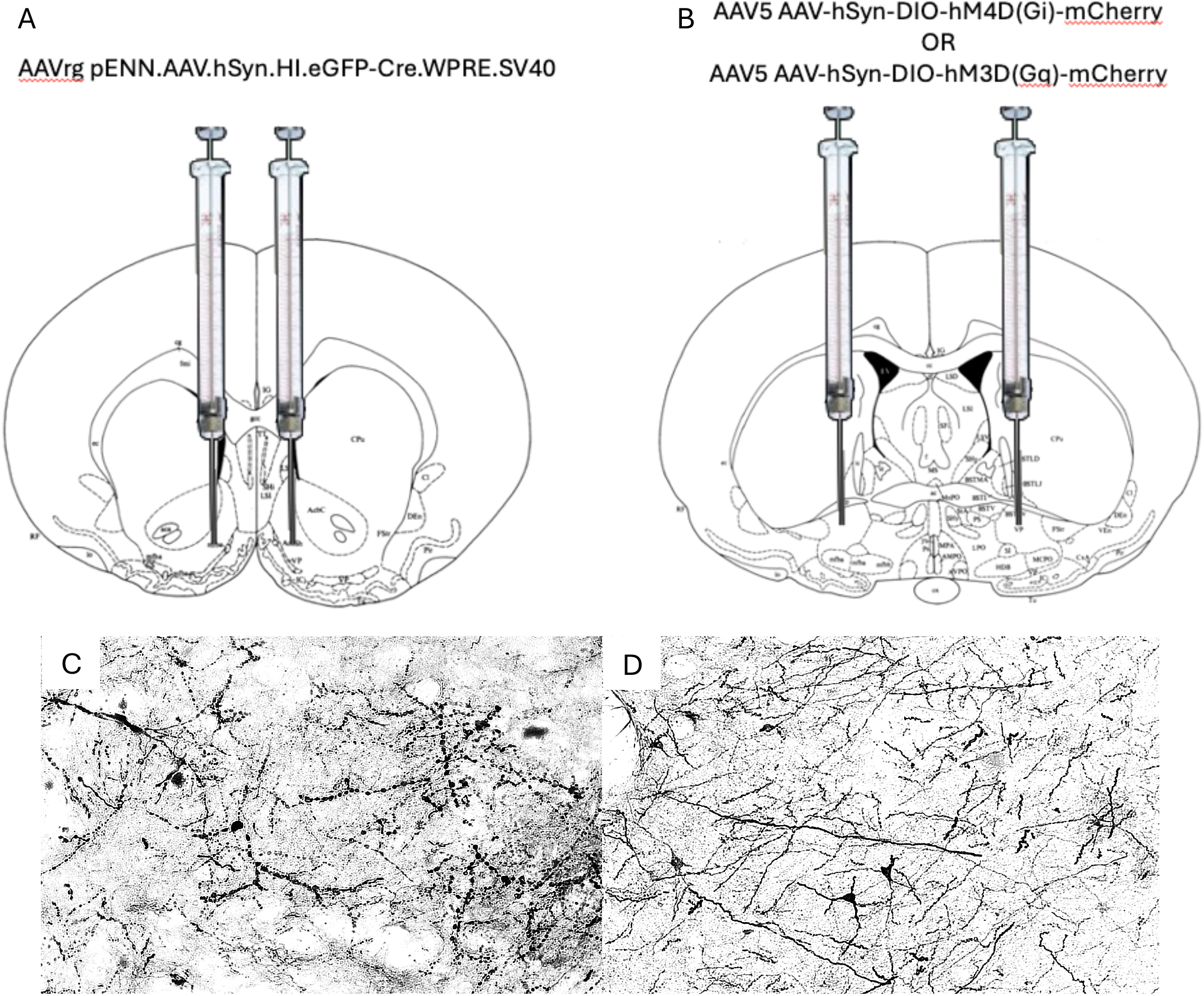
A. A retrograde AAV-Cre viral vector was injected into the AcbSh. B. An AAV DIO construct containing an inverted form of either Gi or Gq DREADD was injected into the VP. Modified from Paxinos & Watson (1986). Representative AcbSh (C) or VP (D) 10x microphotograph of mCherry immunohistochemistry.

## Results

A two-way ANOVA was performed to evaluate the effects of DREADD (excitatory, inhibitory, or no DREADD) and drug administered (vehicle or CNO) on lever presses in a progressive ratio task.

The results indicated no significant main effect for DREADD, F(2, 31) = 2.421, p = .105; no significant main effect for drug administered, F(1, 31) = 2.004, p = .167; and no significant interaction between DREADD and drug administered, F(2, 31) = 1.780, p = .185 (figure 2 A).

**Figure 2.**
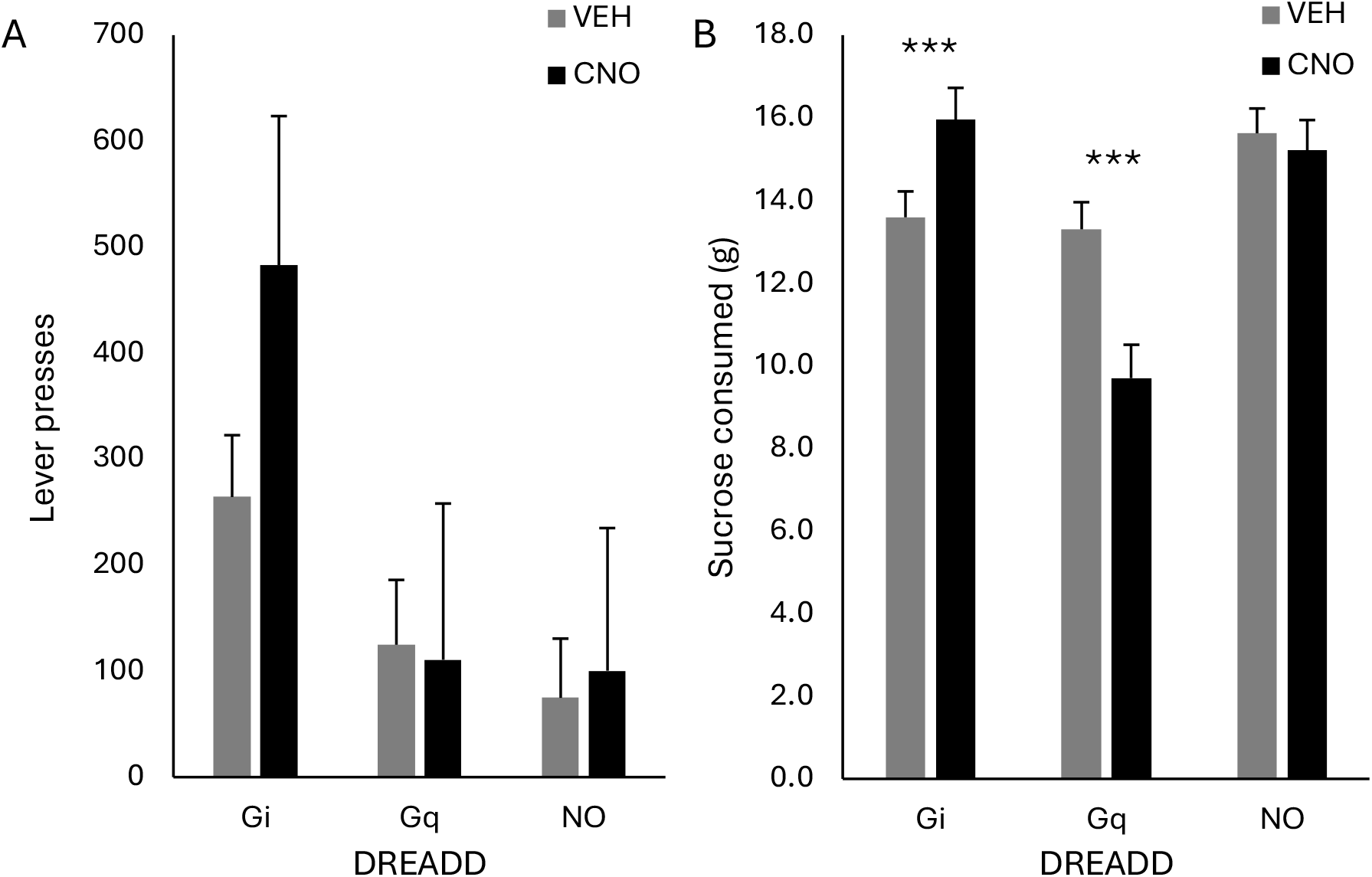
A. CNO administration did not affect motivation to work for sucrose as measured using a progressive ratio task in non-food-deprived DREADD expressing rats (Gi and Gq) nor control rats (NO) (mean ± SEM). B. Non-food deprived rats expressing inhibitory (Gi), excitatory (Gq), or no DREADD (NO) were given one hour to consume 20% sucrose solution after being injected with either vehicle or CNO. CNO-induced chemogenetic inhibition of VP-AcbSh pathway increased sucrose consumption in rats (p = 0.001), excitation decreased it (p = 0.001) and had no effect on rats not expressing DREADD (p = 0.5) (mean ± SEM).

A two-way ANOVA was performed to evaluate the effects of DREADD (excitatory, inhibitory, or no DREADD) and drug administered (vehicle or CNO) on 20% sucrose consumption in non-food-deprived rats.

The results indicated a significant main effect for DREADD, F(2, 31) = 11.170, p = .001; no significant main effect for drug administered, F(1, 31) = 3.148, p = .086; and a significant interaction between DREADD and drug administered, F(2, 31) = 18.891, p = .001 (figure 2B).

Post hoc testing using Bonferroni correction for multiple comparisons indicated that sucrose consumption was significantly higher for rats expressing inhibitory DREADD when CNO was administered than when vehicle was administered (p = .003). Additionally, sucrose consumption was significantly lower for rats expressing excitatory DREADD when CNO was administered than when vehicle was administered (p = 0.001). There was no significant difference between the sucrose consumption of rats expressing no DREADD with administered either CNO or vehicle (p = .5) (figure 2B).

## Discussion

In female rats, chemogenetic excitation or inhibition of projections from the VP to the AcbSh influenced consumption of a 20% sucrose solution but had no effect on the motivation to work for food as measured using a progressive ratio task. Specifically, chemogenetic activation of projections from the VP to the AcbSh in non-food-deprived female rats led to a decrease in 20% sucrose consumption. Conversely, chemogenetic inhibition of the same projection increased consumption of the 20% sucrose solution.

In contrast, Scott et al. (2024) reported that chemogenetic activation of VP projection neurons resulted in no significant changes in food consumption. This apparent discrepancy between the two studies can be explained by multiple reasons. Possibly the most crucial difference between both studies is that here we used a dual vector approach to express DREADD in VP neurons that project to the AcbSh, while Scott et al. (2024) used a single vector approach leading to all GABAergic VP projection neurons expressing DREADD.

Thus, here chemogenetic manipulations affected a small subset of VP projection neurons, namely those that project to the AcbSh, while in the study conducted by Scott et al. (2024), all VP projections were affected by chemogenetic modulation. It is nonetheless informative that we observed different behavioral effects, as this suggests that different VP efferents might have a variety of behavioral effects. This matter could be addressed by future studies dissecting the role of each VP efferent. Additional studies should also consider the sex differences noted by Scott et al. (2024).

Other differences to consider between both studies include the concentration of sucrose, as we used a 20% concentration while Scott et al. (2024) used 10%, the fact that our rats remained pair-housed and opposed to single-housed, and differences in rat strain as they used Long Evans and we used Sprague-Dawley rats. Additionally, there were also differences in the DREADD agonist used, JHU37160 versus CNO in our experiment. While all these differences likely contributed to some extent to the different behavioral results between both studies, we consider that the most likely difference stems from targeting all GABAergic VP projecting neurons in Scott et al. (2024) versus VP neurons projecting to the AcbSh as done in this study.

The directionality of the VP-AcbSh pathway has also been studied by Smedley et al. (2019). Interestingly, this group saw no effect on free feeding on male rats when the projections from the VP to the AcbSh were chemogenetically inhibited. Besides the sex differences in the subjects, it is also notable that Smedley et al. (2020) measured the intake of standard rat chow. In contrast, here we measured the consumption of a 20% sucrose solution. It is then possible that either or both factors, sex and food stuM, might contribute to the different behavioral results observed. Thus, it appears that projections from the VP to the AcbSh mediate food consumption but not motivation to work for food. Future studies looking at other VP effects might be able to dissect which projections are involved in the motivation to work for food.

Additionally, it has been reported that pharmacological activation of the VP leads to increased preference for fat consumption (Covelo et al., 2014). In contrast, the food used in this study contained mainly carbohydrates. Future studies should consider the possibility that different behavioral effects might be observed using fats or offering a choice of different macronutrients.

Further, VP GABAergic neurons send an inhibitory input to the AcbSh, including a subpopulation of ventral arcopallidal neurons (Vanchez et al., 2020). In the study, activation of the VP-AcbSh pathway led to an increase in the consumption of a palatable food reward (Vanchez et al., 2020). In contrast, in our study, activation of the VP-AcbSh pathway led to a decrease in the consumption of the food reward. This discrepancy could be caused by the difference in the nature of the projection neurons recruited, as we targeted all VP neurons projecting to the AcbSh, and not specific subpopulations. It is then possible that the behavioral effects of the whole VP-AcbSh pathway differ from that of specific neural subpopulations.

In conclusion, our findings indicate that the VP-AcbSh pathway mediates the consumption of a palatable sucrose solution. Chemogenetic manipulation of VP projections to the AcbSh selectively influenced sucrose intake without affecting motivation to work for food, suggesting that distinct VP efferents play differential roles in feeding behavior versus food-seeking motivation. The discrepancies observed across studies of this pathway, both in terms of projection targets and specific neuronal populations. Additionally, the findings indicate a nuanced role for the VP-AcbSh pathway in modulating the intake of specific macronutrients. Future studies that dissect the role of the VP-AcbSh pathway should consider the variables such as food type, sex, and neural subpopulations as well as their interactions.

## Acknowledgments

The authors would like to thank the College of Natural and Health Sciences for the support, the NIDA – Drug supply program for providing CNO, and the following students for their help with this work: Megane Beaupre, Ari Hjelmeseth, Miranda Johnson, Hedi Morris, Zach Wilson, Aryan Patel, Mark Huckeby, and Dylan Jensen.

## Notes

### Competing Interest Statement

The authors have declared no competing interest.

